# Putting the brain in a box: a toolbox for creating Cartesian Geometric Representations with Isometric Dimensions (Cgrids)

**DOI:** 10.1101/786368

**Authors:** Mathijs Raemaekers, Mark Bruurmijn, Nick Ramsey

## Abstract

The folding of the human cortex complicates extraction of position information and recognition of patterns across the cortical surface. As straight lines correspond better to our intuitions in spatial orientation, we developed an approach for imposing Cartesian grids on portions of the cortical surface, which can then be represented in a rectangular matrix. These functions have been implemented in the Cgrid (Cartesian Geometric Representation with Isometric Dimensions) toolbox. Cgrids can be generated based on regions of interest, or combinations thereof, according to any one of the Freesurfer’s annotation schemes.

The toolbox was evaluated using the surface reconstructions of T1-weighted images of 30 subjects, and 17 different Cgrids that in combination covered nearly the entire surface area of the brain. The vast majority of Cgrids (94.6%) could be generated without issues.

In addition to facilitating spatial orientation and pattern recognition, the toolbox also allows detailed comparison between the left and right hemisphere, and performing surface based analysis using a volumetric data format. The output of the toolbox is fully compatible with most existing fMRI/MRI analyses packages, and is immediately suitable as input for second level analysis. The toolbox can be downloaded from: https://github.com/mathijsraemaekers/Cgrid-toolbox

## Introduction

The human cortex is composed of many gyri and sulci, which result from its heavy folding/gyrification. Although this folding is an evolutionary necessity, allowing the brain to fit inside a relatively small cranium with increased size of the cortical sheath while simultaneously reducing neuronal wiring length (Welker, 1990), its geometric features do not connect very well with our human capabilities in spatial localization. Orienting through the brains curved gyri and sulci can be a difficult endeavor, like finding your way in medieval European city centers. Surface reconstructions (Dale et al., 1999) and subsequent steps such as inflation and flattening can most certainly help matters (Fischl et al., 1999), but even after these steps it’s still difficult to deduce readily interpretable positioning information.

In the current manuscript we describe and evaluate an approach for imposing Cartesian grids on portions of the cortical surface, which can then be represented in a rectangular shape. The approach represents a generalization of the methods proposed by Bruurmijn et al., which applied similar techniques to the sensorimotor cortex (Bruurmijn et al., 2019). This procedure can substantially facilitate the interpretation of spatial patterns of anatomical or functional features, and produce straightforward metrics of location. The necessary processing steps have been implemented in the Cgrid (Cartesian Geometric Representation with Isometric Dimensions) toolbox, which is an extension to the FreeSurfer software package, and uses the output of the FreeSurfer “recon-all” pipeline as input (Dale et al., 1999; Fischl et al., 1999). The Cgrid-toolbox provides a user friendly way to design and generate Cartesian grids of functional and anatomical data within predefined portions of the cortical surface. The output of the toolbox is fully compliant with all the major MRI/fMRI analysis and visualization tools.

## Overview of the Procedure

Figure 1 provides a pictorial overview of the main processing steps. Cgrids can be generated for any portion of the cortical surface for which 4 borders can be defined (upper, lower, left, right) based on any one of the FreeSurfer annotation schemes. Cgrids can also be composed of multiple adjacent subgrids, for each of which 4 the borders must be defined independently. Each subgrid border is composed of one or a combination of borders between FreeSurfer ROI-pairs. When a Cgrid is composed of multiple subgrids, they are vertically concatenated, and thus must share their upper/lower border. The surface area enclosed by the borders is subdivided over the vertical axis by first fitting 2 high-order polynomials, one through the upper border and one through the coordinates of the lower border. Then intermediate polynomials are generated, one for each vertical resolution element required in the end-result, by linear interpolation of the polynomial coefficients of the models of the upper and lower border. Subdivision over the horizontal axis is achieved by subdividing the polynomial models into portions of equal length, or by repeating the polynomial modeling and interpolation procedure for the left and right border. The fitting and subdivision is repeated for every subgrid in the Cgrid. After the procedure, functional and anatomical information can be mapped onto the Cartesian grid, and stored in nifti format. The procedure is implemented in the Interactive Data Language (Harris Geospatial Solutions, Colorado, USA) and is described in full detail here below.

**Figure 1.**
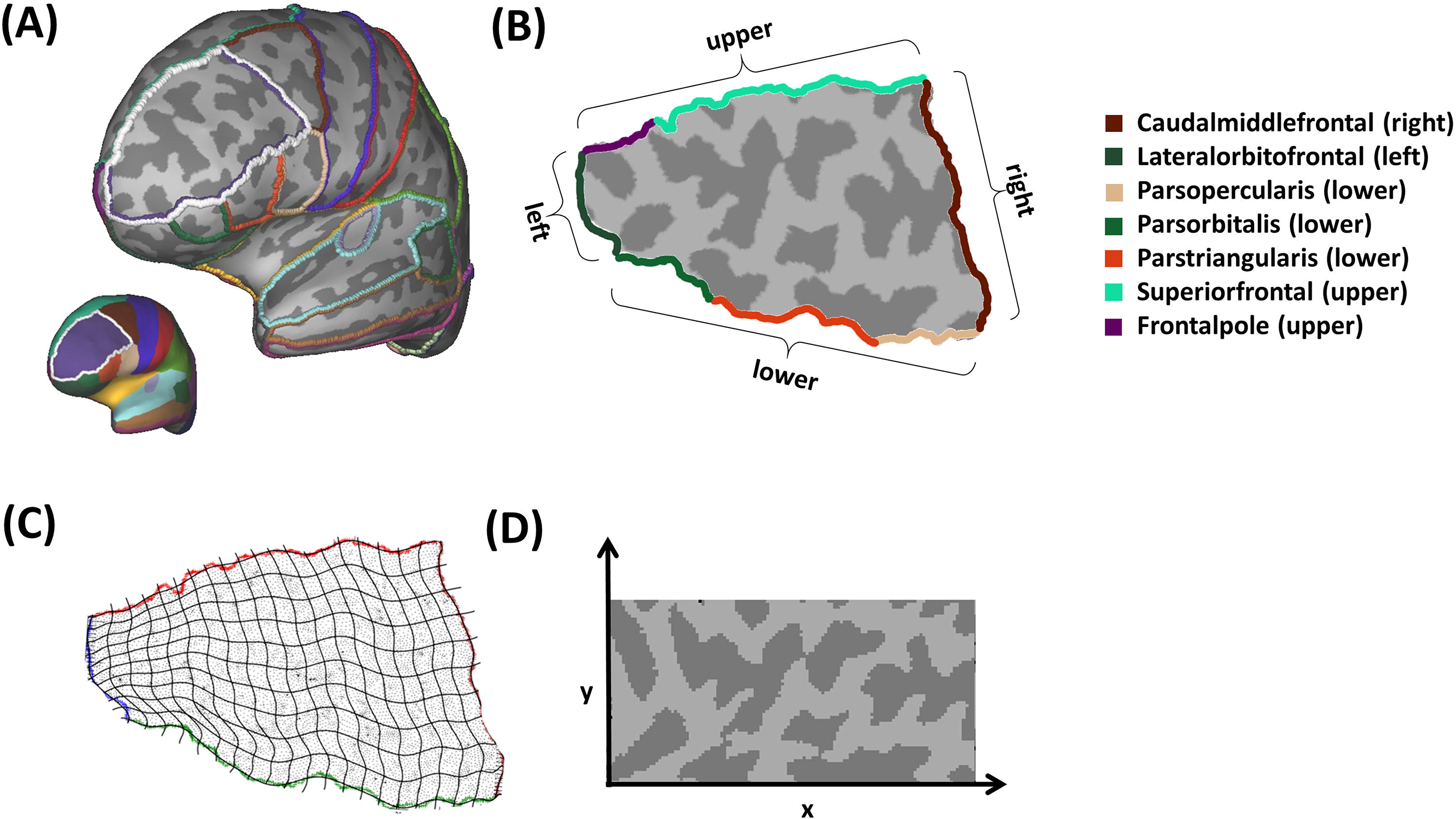
Basic overview of the procedure for creating Cgrids. (A) A portion of the cortical surface is selected based on the labels of available annotation schemes. For this example we chose the rostral middle frontal cortex (encircled in white), as defined by the DK atlas; (B) The patch is extracted and subsequently flattened, and 4 borders are defined based on the borders between one or multiple ROI pairs; (C) High order polynomials are fitted through the upper and lower border, followed by the generation of intermediate interpolated polynomials, effectively construing the Cgrid y-coordinates for each vertex. Cgrid x-coordinates of nodes are established by subdividing the polynomials into portions of equal length, or by repeating the polynomial modeling and interpolation procedure for the left and right border. Note that the number of visualized interpolated polynomials is reduced for clarity; (D) The Cgrid x and y coordinates of each vertex can then be used to map surface based information to rectangular matrices, which can be stored in nifti format.

### 1. Generating patches

The first step in the procedure is the extraction of patches, which are portions of the surface reconstruction on which the Cgrid is to be applied. Patches include the nodes of a single or a combination of adjacent ROIs from one of the FreeSurfer parcellations, and should minimally encompass the entire cortical surface area on which the grid is to be imposed. Patches are “dilated” to also include the first order neighboring nodes of the selected ROIs. These patches are extracted from the inflated surface, stored as FreeSurfer patch file, and subsequently flattened using FreeSurfer’s “mris_flatten” algorithm (Fischl et al., 1999), with the option to set the flattening parameters for reducing either processing time or metric distortions.

### 2. Defining Borders

Cgrids are composed of one or multiple subgrids, and four borders need to be specified for each subgrid, i.e. an upper, lower, left, and right border. Borders are sets of coordinates that represent the middle of the edges linking nodes of 2 different FreeSurfer ROI’s in the flattened patch. Each subgrid border can be specified as one or a combination of borders between 2 FreeSurfer ROIs. The combination of FreeSurfer ROI borders that define the different subgrids are stored in a Cgrid-scheme, that includes the parameter settings of a particular Cgrid, which can be applied to the surface data of multiple individuals.

Coordinates or clusters of coordinates can be optionally removed from the border if their distance to the main cluster exceeds a specified threshold, which allows negating errors in the cortical labeling. Borders (upper, lower, left, right) are always defined based on the perspective within the left hemisphere. For the right hemisphere, left and right borders are swapped, and the horizontal-axis reversed. This is to ensure that Cgrids of the left and right hemisphere are homotopically mapped onto each other.

### 3. Modeling the upper and lower border

In an initial re-orientation step using straight line fitting with errors on both coordinates (Press et al., 2007), the flattened patches and their borders are rotated in such a way, that the upper and lower borders run maximally parallel to the horizontal axis. When after rotation the upper border is located below the lower border, the patch and borders are rotated an additional 180 degrees to correct for this.

The next step includes the generation of high-order polynomial models for the upper and the lower border. The polynomial models are optimized by minimizing the averaged (across upper and lower border) root mean square of the deviation between the predicted and real y-coordinates of the border. The optimization includes simultaneously:

1: Rotating the patch; there are limitations to what the high order polynomials can accurately model, including high frequency spatial oscillations in the shape of the border, or the presence of multiple y-coordinates for a single x-coordinate. The influence of these limitations is minimized by applying an optimized additional rotation step (search between −90° to 90° in steps of 1°)
2: Changing the polynomial order; while in principle higher order polynomials should produce models with less error, we observed breakdowns in the fitting procedure for polynomials with order ⍰ 15, where the error of the model increased with higher orders. We speculate that this might be due to round off errors, as the coefficients for the high order variables become increasingly small. The optimal polynomial order was chosen, by estimating the amount of error in the fit for all polynomial models between order 1 and 30
3: Extrapolating the upper and lower borders; in subsequent steps the fitted polynomials are used to establish Cgrid y-coordinates along the entire horizontal range of the patch. However, for high order polynomials, the shape of the model becomes highly volatile immediately beyond the x-range of the upper or lower border coordinates. Patch x-coordinates can however exceed the x-range of the upper and lower borders, and for such coordinates the established polynomial model is inappropriate. This issue is solved by extrapolating the upper and lower border, so that they cover the entire horizontal range of the patch. This extrapolation involves adding coordinates to the border’s endpoints at the same spatial density per horizontal mm as the rest of the border, so that extrapolated portions are weighted equally in the polynomial fit. Coordinates are added in such a way, that the slope of the extrapolated part gradually changes from the slope at the endpoint of the border (as estimated based on the last 10 coordinates of the border), to a slope that is approximately perpendicular to the left or right border. These perpendicular slopes are established by fitting straight lines through the upper and lower half of the left and right border(Press et al., 2007), and subsequently taking one over the slope of the calculated fits. Figure 2 shows the border extensions for the Cgrid of the Rostral Middle Frontal cortex, that was depicted in Figure 1. Border extensions start of running parallel to the border endpoints, and gradually change slope until they run approximately perpendicular to the left and right borders at the extremities of the x-range of the patch.

**Figure 2.**
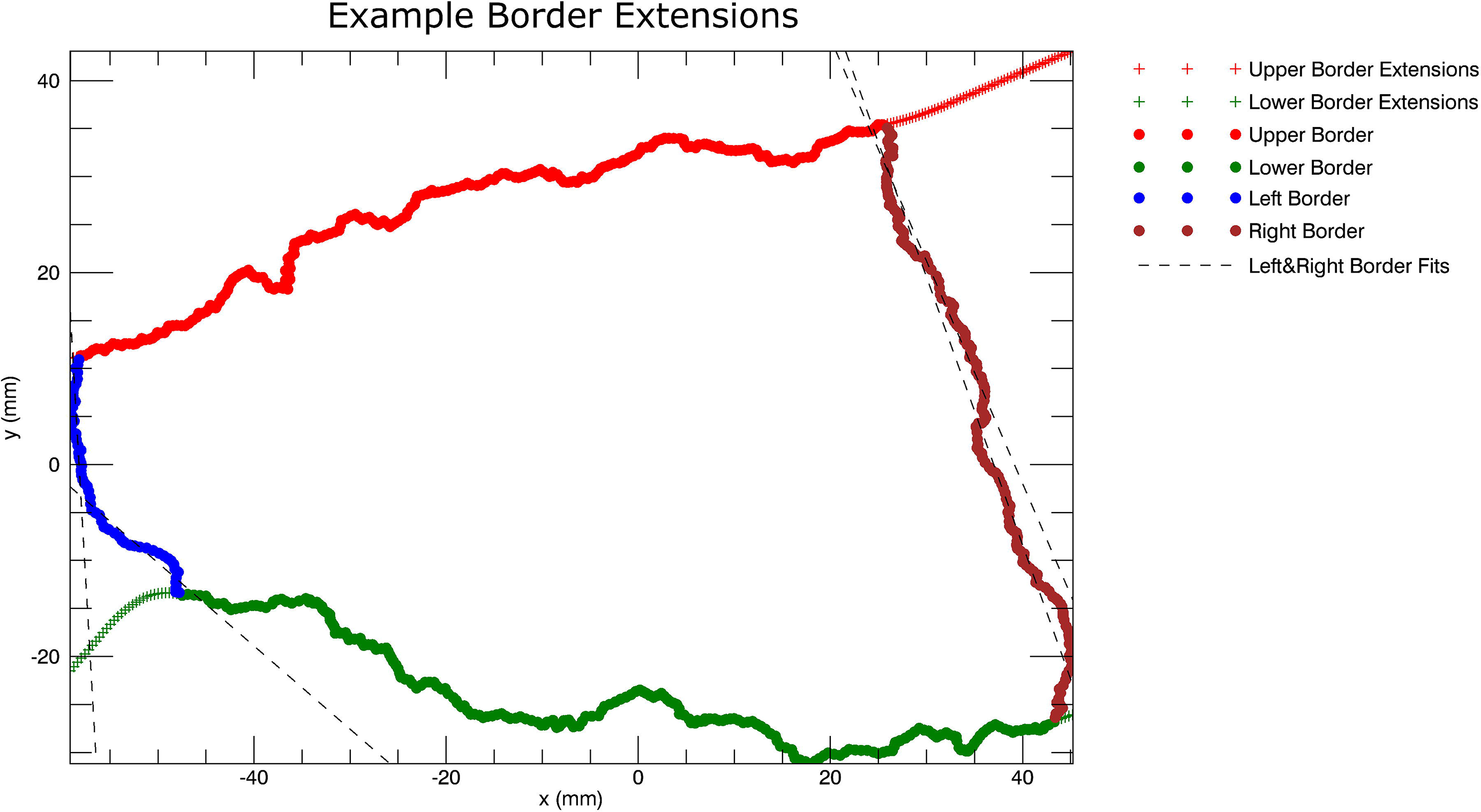
Visualization of the procedure for the upper and lower border extensions. The upper and lower borders need to be extrapolated to extend the full horizontal range covered by the nodes to be included in the subgrid. Straight lines are fitted through the upper and lower half of the left and right border (dashed lines, 4 in total), which are then used to calculate the slope perpendicular to the upper/lower halves of the left and right border. Upper/lower border extensions (plus-signs) start with the same slope as the end portions (i.e. the last 10 coordinates) of the upper and lower borders. The slope then gradually changes until extensions run perpendicularly to the upper and lower halves of the left and right borders, so that interpolated polynomials will cross the left and right borders approximately perpendicularly.

### 4. Establishing Cgrid y-coordinates

The polynomial modeling results in 2 sets with an equal number of parameters, one for the upper and one for the lower border model. Intermediate polynomial models are generated by (for each polynomial order) linearly interpolating between the parameters of the models for the upper and lower border, with the number of interpolation steps determining the number of tiles/steps along the vertical dimension. Subsequently, the predicted y-coordinate is calculated for the x-coordinate of every node in the patch, and for all polynomial models (both those fitted to the upper and lower border and those interpolated between the borders). The Cgrid y-coordinate is determined for each node by establishing if the real y-coordinate of a node is larger or smaller than each of the predicted y-coordinates, and establishing between which adjacent polynomial models the node is located.

### 5. Establishing Cgrid-x-coordinates

The toolbox offers two approaches to establishing Cgrid x-coordinates for the nodes in the patch.

1. 2-polynomial approach: Polynomial curves are drawn based on the models generated in the previous step and extend the entire x-range of the patch, using a step size of 0.1 mm. For the resulting set of coordinates of each polynomial curve it is established which coordinates are closest to any coordinate of the left and the right border, and these are marked as start/end-point. This step can optionally be preceded by spatially smoothing the coordinates of the left and right border with a Gaussian kernel, to reduce high frequency spatial oscillations that can result in multiple crosses between the left/right borders and the polynomial curves. The total length of the polynomial curves between start-points and end-points is calculated, and markers are placed at equidistant locations along the curves. The number of markers is determined by the desired x-resolution of the Cgrid. Subsequently, the x-coordinate for each node in the patch is established by assessing if it is located left or right of straight lines running between equivalent markers of the adjacent polynomials between which the node is located.
2. 4-polynomial approach: The second approach effectively repeats the procedure from the initial reorientation of the patch onwards, but the definitions of upper and lower borders are swapped with those of the left and right borders. This rotates the patch approximately 90°, and then performs the polynomial modeling on the left and right border. This is similarly followed by the interpolation if the polynomial models, and subsequent establishment of the Cgrid x-coordinates of the nodes in the patch.

When the Cgrids is composed of multiple subgrids, they are vertically concatenated, so that the y-axis runs continuously across the subgrid. Using the 2-polynomial approach will avoid spatial shifts along the upper/lower border where the subgrids are concatenated, as equidistant locations of the polynomial models are closely matching for the upper border in one subgrid, and the lower border in an adjacent subgrid. This approach is thus advantageous when Cgrids are composed of multiple subgrids, as it allows proper concatenation of subgrids along the x-axis. The 4-polynomial approach is generally more straightforward, as it treats the borders along the two axes similarly. For this approach however, the Cgrids do not run continuously across subgrids, and should thus be used only when Cgrids include a single subgrid. Results of the 2 and 4 polynomial approach for the area depicted in figure 1 can be seen in figure 3.

**Figure 3.**
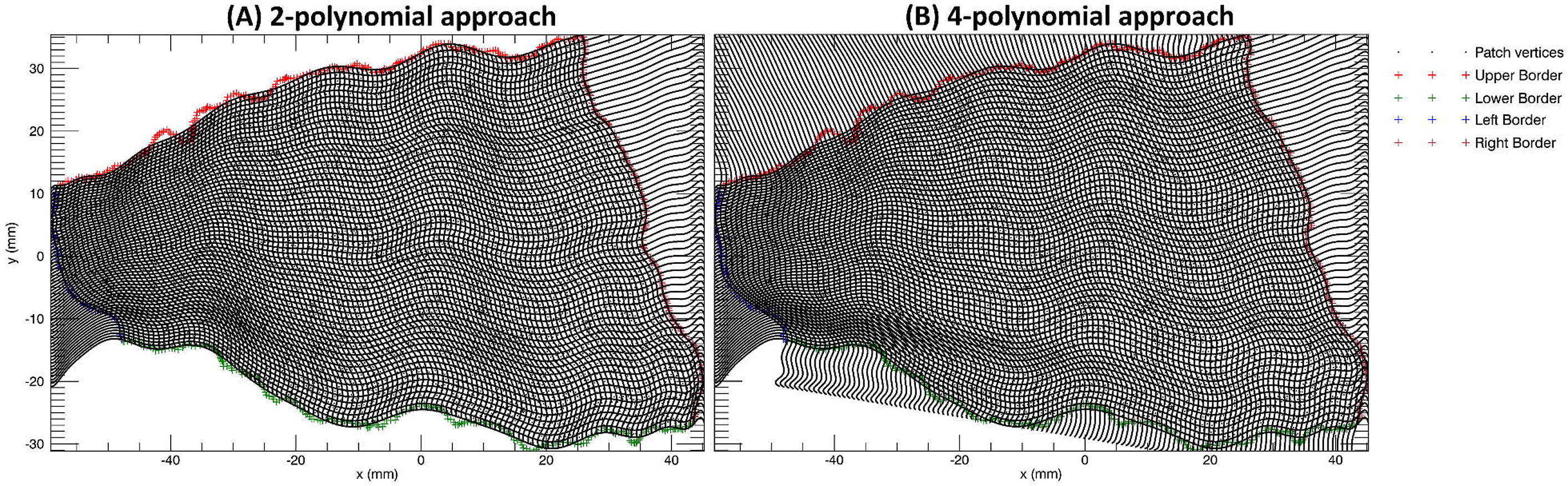
120 × 60 Cgrids imposed on the left rostral middle frontal ROI as the one depicted in figure 1. Both panels show the polynomials fitted through the upper and lower border coordinates, in addition to the 59 intermediate polynomials that are generated by linearly interpolating the parameter estimates. (A) Represents the 2-polynomial procedure, where the x-coordinates are determined by drawing lines between equidistant locations subdividing the polynomials in 120 portions of equal length, and drawing lines between equidistant locations on interpolated polynomial (B) Represents the 4-polynomial approach where the patch is rotated 90 degrees and the fitting procedure is repeated along the second axis.

The result of the described procedure is a matrix including the vertex-number of all nodes within the Cgrid, in addition to their Cgrid x and y-coordinates. At this stage, the x and y-coordinates can be swapped and the x-coordinates reversed to allow full flexibility in the orientation of the axes of the final Cgrid. The Cgrid-coordinates for each node are used to generate the different outputs of the toolbox. In addition, the toolbox includes various options to monitor the procedure, including the provision of images as those depicted in figure 2 and 3, that can be used to evaluate the quality of the results, and to identify potential causes of errors.

## Evaluation of the procedure

For the evaluation of the procedure we used the T1-weighted images of 30 subjects that participated in a previous study. All subjects gave approval for the usage of their data. Parameters of the T1-weighted image sequence included: TR 8.4 ms, TE 3.8 ms, flip angle=8°, FOV 288×288×175, matrix 288×288×175, voxel size 1.0×1.0×2.0 mm, 175 slices, sagittal orientation, total scan duration 487 seconds. T1 weighted images were processed using the standard recon-all pipeline of FreeSurfer 6.0.

17 Cgrid-schemes were drafted that in combination covered nearly the entire surface of the cortex. The 17 Cgrids contained 27 subgrids, and were generated for all 30 subjects. The Cgrids and their properties can be seen in the 7 leftmost columns of table 1 and in figure 4. The complete schemes as used here can be downloaded from the toolbox’s website. The schemes were all based on the parcellation according to the DK-atlas (Desikan et al., 2006), except for the language Cgrid which was based on the DKT-atlas (Klein and Tourville, 2012). The DKT atlas was chosen for the language Cgrid, as the parsorbitalis has only 3 borders in the DK-atlas, but four in the DKT atlas, as required by the toolbox.

**Figure 4.**
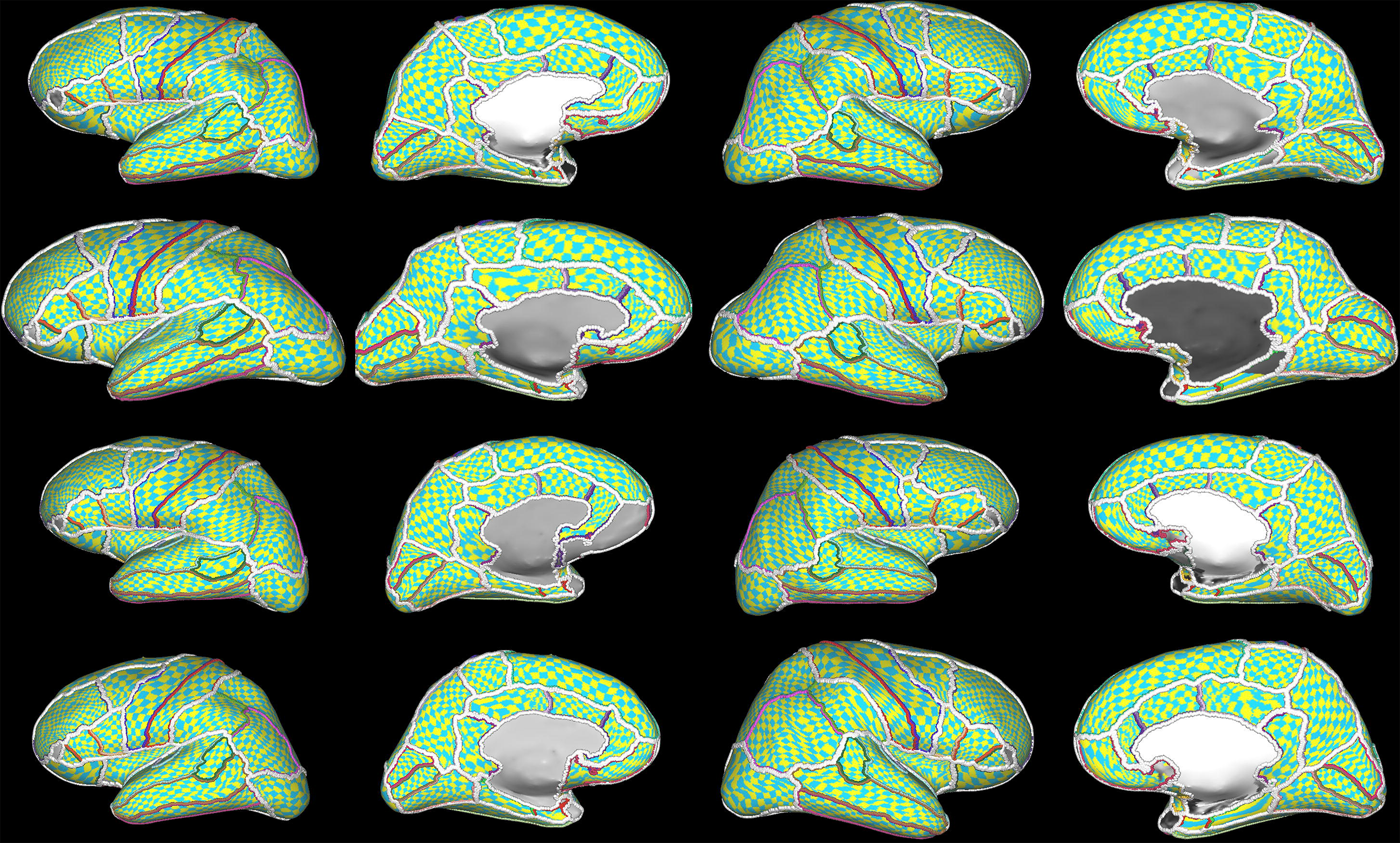
The coverage of the 17 Cgrids of 4 included subjects projected on an inflated surface. Each row is of a different subject, showing the lateral and medial views of the left and right hemisphere. The white borders run between the different Cgrids. Other colors signify borders of FreeSurfer ROIs. The resolution of displayed tiles was reduced to improve visualization. Note that there is a slight anomaly visible near the pars orbitalis of the language Cgrid, due to the choice of a different annotation (DKT-atlas) for the language Cgrid as opposed to the other Cgrids (DK-atlas). The algorithm failed for the left medial orbitofrontal Cgrid of the third subject.

The results were appraised both by estimating the amount of error in the polynomial fitting procedure and by visual inspection of the graphical output of the toolbox. Both the mean and maximum deviation between the border coordinates and the polynomial model were stored and evaluated, with the maximum deviation used as metric to identify to presence of large local errors in the fitting procedure. Using visual inspection, we classified 5 types of errors in the generation of the Cgrids that can significantly reduce performance. While some of these errors can be fixed by additional processing steps or alternative choices in the composition of the Cgrid-schemes, we abstained from doing so to illustrate limitations and caveats in the use of the toolbox. Types of errors included:

1) Erroneous border extrapolation: The upper and lower borders are extrapolated so that they run approximately perpendicular to the left and right border. However, the left and right border can be shaped in such a way that running perpendicular in the direction of the extrapolation means cutting through the patch. These errors can be detected by visual inspection of the toolbox’s graphical output.
2) Not all borders detected or borders consist of too few coordinates: The existence of some ROI borders varies across subjects, so that for particular Cgrids in some subjects some subgrid borders cannot be defined. Note that the absence of single ROI border need not be problematic when a subgrid border consists of multiple ROI-borders. All subgrid borders require a minimum of 3 coordinates, to be useful. This type of error is logged by the toolbox.
3) Crossovers: The projection during flattening can result in portions of the surface crossing over each other, causing topological defects in the flattened patch. These translate into errors in the Cgrid and potentially subgrid borders crossing over on itself. This type of error can be detected by visual inspection of the toolbox’s graphical output, and also increases the amount of error in the polynomial fitting procedure. An example of this type of error is shown if figure 5B.
4) Islands: When not all coordinates in the patch are interconnected, this can result in multiple independent border sections between 2 ROIs. The toolbox includes algorithms for removing these islands of border coordinates, but these are currently not 100% effective as they can remove unintended parts from borders. This type of error can be detected by visual inspection, and also increases the amount of error in the polynomial fitting procedure.
5) Interpolated polynomials crossing the left or right border at multiple locations (2-polynomial approach only), causing possible errors in the estimated start/end-locations for the subdivision of the polynomials in equidistant portions. These errors can be detected by visual inspection of the graphical output of the toolbox. An example is shown if figure 5C.

**Figure 5.**
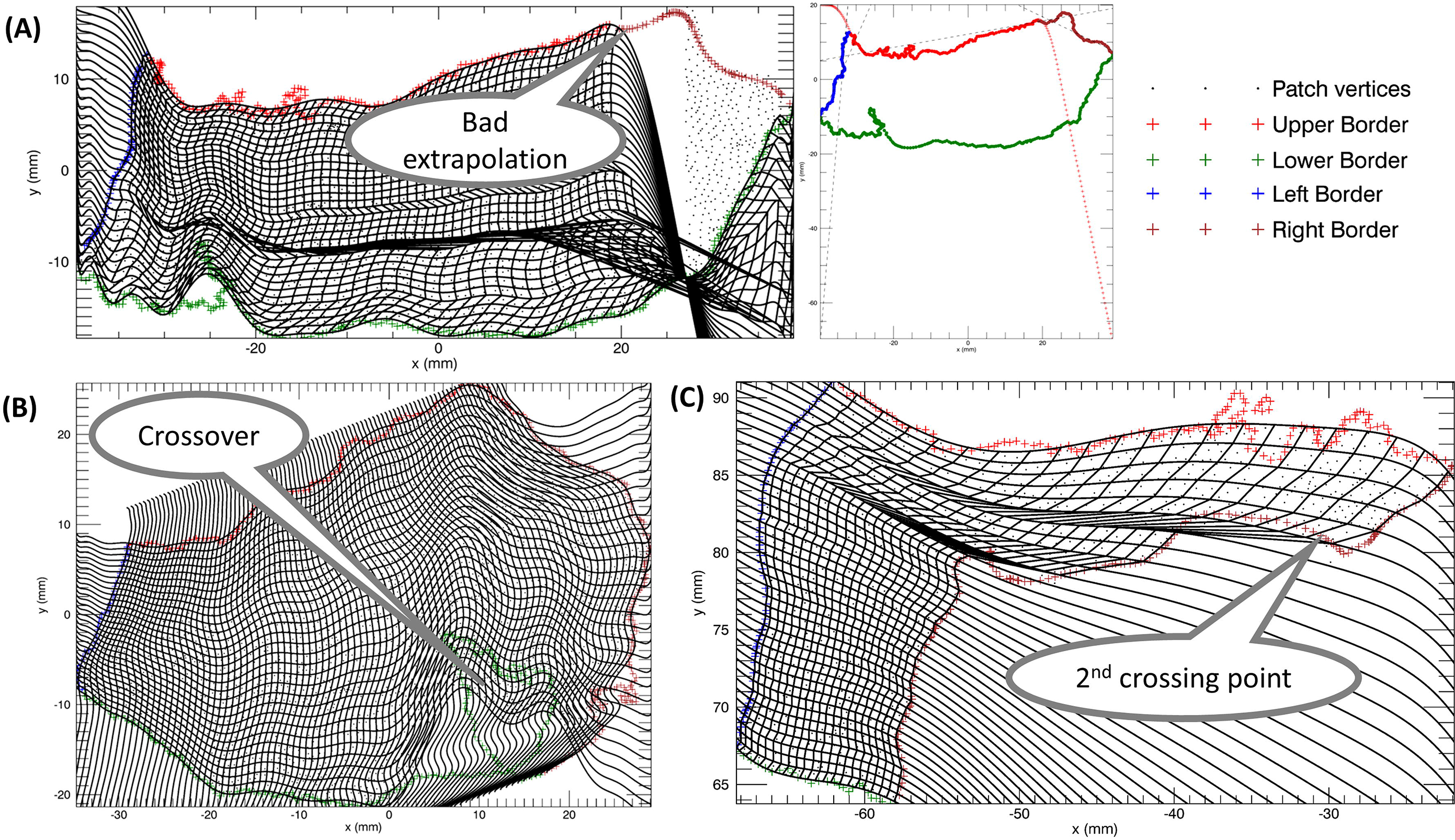
Typical examples of the most frequently encountered types of errors when running the procedure. The figure shows plots like those presented in figure 3&4. (A) Example of of an erroneous extrapolation of the right upper border of a medial orbitofrontal ROI. The border is extrapolated to run perpendicular to the upper part of the right border. Here the perpendicular direction (with increasing x-coordinate) means cutting through the patch. (B) A crossover in the lower border of an insula subgrid results in multiple y-coordinates for a single x-coordinate, and subsequent error in the polynomial fit. (C) Example of multiple crossings between an interpolated polynomial (5 polynomial from the top) and the right border of a rostral anterior cingulate subgrid. As start/end-points of polynomial models cannot be determined unequivocally, the determination of the Cgrid x-coordinate is compromised

## Results of the evaluation

The vast majority of all subgrids were generated without one of the described errors (1533/1620=94.63%; 95% confidence interval 93.42%−95.68%). The mean square error of the polynomial fits, averaged across all subgrids, was 0.44 mm, and the mean maximum deviation was 2.30 mm. Results per Cgrid can be seen in the 8 rightmost columns of table 1.

Of the total of 87 erroneously generated subgrids, 76 (87%) were of the insula, parahippocampal+entorhinal, rostralanteriorcingulate or medialorbitofrontal ROIs. Excluding subgrids from these ROIs raised the success rate to 99.2% (95% confidence interval 98.58%−999.60%). Performance for most subgrids was thus near perfect, while a limited number of schemes encountered issues on a regular basis.

Difficulties in the insula were caused by frequent and relatively large crossovers after flattening, and thus fitting errors. Increasing the number of iterations during inflation of the surface can in all likelihood reduce the number and extend of the crossovers, but this has to be done separately after the FreeSurfer recon-all pipeline.

For the parahippicampal+entorhinal and the rostral anterior cingulate Cgrids, the majority of errors involved multiple crossing points between the left/right borders and interpolated polynomials, which resulted from their particular shape and constellation of borders. The most prominent issue with the medial orbitofrontal Cgrid was incorrect extrapolation of the borders, caused essentially by the shape of patch as defined by the borders too much deviating from a rectangle. Some of these issues might be solved by more extensive border smoothing, but also the scheme of the Cgrid for covering a particular part of the cortex might be reconsidered, e.g. by using the DKT or the a2009s atlas instead of the DK atlas.

## The output of the toolbox

The labeling with Cartesian x and y-coordinates of nodes within the Cgrid can be used to create various output.

Surface data can be mapped to the grid by combining the values from nodes within tiles. The data can then be stored in Cgrid nifti-format, which is a regular nifti file of which the first two dimensions represent the Cgrid x and y coordinates, and the third dimension reflects the left and right hemisphere. The fourth dimension can optionally be used for timeseries data.

Various surface based metrics based on the FreeSurfer ‘recon-all’ pipeline can be mapped to the Cgrid nifti format. These include for each Cgrid tile (1) the number of nodes (2) total white/pial surface area (3) total grey matter volume (4) mean cortical thickness (5) mean cortical depth and (6) mean curvature. An example is shown in figure 6A. In addition, volumetric MRI/fMRI data can be transformed to the Cgrid nifti format, where volumes are first mapped to the surface using any one of the algorithms provided by FreeSurfer’s “mri_vol2surf”. When averaging values of nodes within Cgrid tiles, they are weighted according to their area across the cortical surface. An example of an fMRI analysis using Cgrid nifti-files is show in figure 6B. Missing data as a result of Cgrid tiles that do not contain any nodes, can optionally be filled in. Tiles can be empty when they are smaller than the internode distance, either because of the choice for a high resolution Cgrids, or because of the shape of the patches on which the subgrids are based. The filling in of missing values is done by creating a copy of the data that is smoothed in two dimensions with a small Gaussian kernel (0.5 pixels FWHM) with the option to ignore missing values, and using the values of the smoothed data to fill in the missing data of the original dataset.

**Figure 6.**
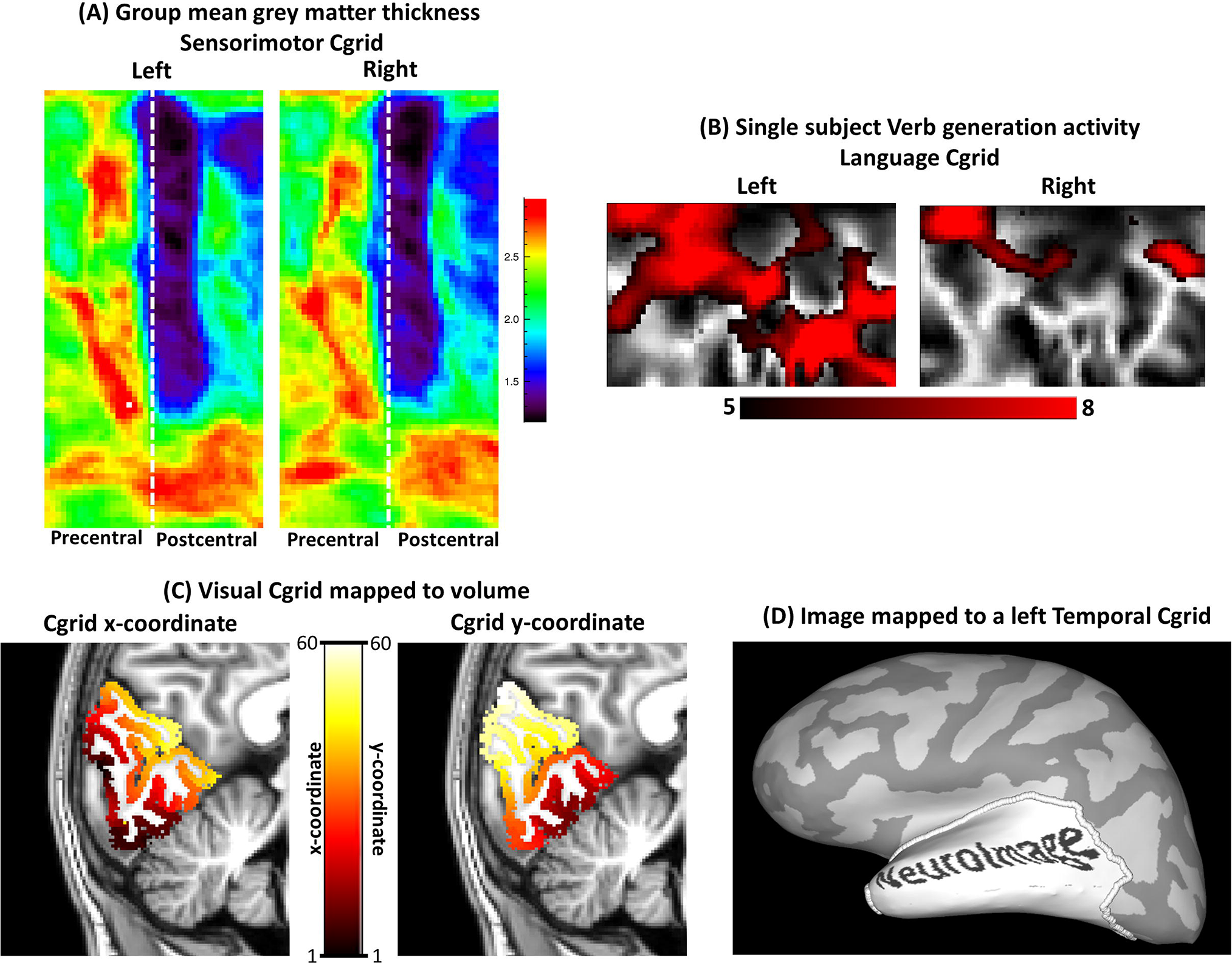
Gallery showing results based on the output of the Cgrid toolbox. Note that the figure is intended primarily for demonstrating optional use of the toolbox. (A) Group mean (n=30) grey matter thickness in the sensorimotor Cgrid. The white dashed line indicates the border between the precentral and postcentral gyrus. (B) Single subject language activity during a verb generation task vs. rest. Functional series were mapped to the language Cgrid with 6 mm surface smoothing and analyzed using multiple regression. Significant pixels are superimposed on curvature data (C) Visual Cgrid mapped to the gray matter in volumetric space. Voxels within the Cgrid are projected on a T1-weighted image and color coded according to their Cgrid x (left) or Cgrid y (right) coordinates.(D) Image mapped to the Cgrid and projected on an inflated surface reconstruction.

This Cgrid nifti format is fully compatibility with most of the main MRI and fMRI analysis software packages and is in principle suitable for second-level analysis, provided that data is appropriately smoothed across the surface during mapping of data to the surface, to account for variations across subjects.

Alternatively, the Cgrid can be mapped to a 3D volumetric space, by labeling voxels with Cgrid coordinates. This results in 2 volumetric nifti volumes for each Cgrid. Each volume contains either the Cgrid x or y coordinates for each voxel within the Cgrid. This procedure uses the Freesurfer algorithm for mapping labels to a volume. While for most applications the Cgrid nifti format is preferable, a volumetric nifti file can be useful in cases where analysis in 2D space is difficult, such as with diffusion data.

Finally, Cgrids can also be used for inversely mapping rectangular images back to the surface of the brain. The result is stored as a FreeSurfer binary surface file, which can be overlaid on a surface representation. These can help to assess the shape of the Cgrid across the surface, or for being used for generating artwork. See figure 6D for an example.

## Discussion

We developed an approach for imposing Cartesian grids on portions of the cortical surface, which is implemented in the Cgrid toolbox, and functions as an add-on to the Freesurfer software package. We drafted Cgrid schemes for 17 areas, which in combination covered nearly the entire surface area of the cortex, and tested these in 30 subjects. For the vast majority of schemes, the toolbox performed near perfectly.

The rectangular representation of portions of the brain can be readily interpreted and positioning information easily deduced. This is especially relevant when function is intimately linked to location within a particular region of interest. Methods for parameterization of location within portions of cortex have been proposed for amongst others the sensorimotor (Coulon et al., 2011) and visual cortex (Benson et al., 2012). The current approach is more general and can be applied throughout the cortex.

An additional property of Cgrids is their inherently defined correspondence between locations in the left and the right hemisphere, so that matching Cgrid-coordinates in the two hemispheres represent homotopic areas. This can be a valuable property for quantifying brain asymmetries, as it allows for direct and detailed assessment of interhemispheric differences in patterns of anatomic or functional features. This can help to investigate previous observed phenomena in more detail (Raemaekers et al., 2018; Zhou et al., 2013), or might be used in the development of more powerful tools for establishing language lateralization (Jones et al., 2011)

The Cgrid toolbox might also help to facilitate surface based analysis in general. While surface based analysis offers several key advantages relative to analysis in volumetric space for a number of applications(Dale et al., 1999), handling of surface based data is often less straightforward, mainly due to compatibility issues across analysis packages and because visualization is more difficult to implement. Although surface based data, such as timeseries data per vertex, can be stored and analyzed in nifti and other formats designed for volumetric data, the format then does not contain any meaningful topographic information and therefore cannot be visualized straightforwardly. The Cgrid nifti format allows surface based data to be compatible with most standard MRI visualization and analysis tools, while maintaining a recognizable topography inside the volumetric file format. In addition, the size of Cgrid nifti files is under most circumstances very small, mitigating storage requirements, and transport and analysis of data.

Cgrids are in a standard space, as particular brain areas of different subjects are morphed into metrically identical Cartesian grids, using ROI borders as defined by automatic cortical automatic labeling (Desikan et al., 2006; Destrieux et al., 2010; Fischl, 2004; Klein and Tourville, 2012) following spherical registration (Bruce Fischl et al., 1999). This implies that the output of the toolbox is suitable as input for second level analysis. The current manuscript does not include a quality assessment of how well this normalization approach performs compared to the possible alternatives, as this may very much depend on the exact scheme for generating the Cgrid and brain area under investigation. However, a previous study (Bruurmijn et al., 2019) found that intersubject alignment of activity during different bodily movements in a Cgrid covering the sensorimotor cortex was comparable to that after spherical registration, and slightly better than for standard volumetric normalization.

The toolbox is designed to be user friendly, flexible, and easily accessible. It requires installment of Freesurfer and the IDL virtual machine, both of which are available free of charge, and the toolbox can be fully driven by a graphical user interface. It supports batch operations for all of its main functions, so that multiple subjects can be analyzed in a single run, facilitating its use in large scale datasets. The toolbox runs fully automatically when making Cgrids based on any one of the cortical annotations that are included in the standard output of the FreeSurfer recon-all pipeline. In addition, Freesurfer enables the possibility to manually edit the existing or generate custom annotations, which can then be used as input for the toolbox. The latter allows a more complete flexibility in defining Cgrids.

We validated the performance of the toolbox using cortical surface reconstructions based on the T1-weighted images of a single 3-Tesla pulse-sequence. Results when using other pulse-sequences and fields strengths might be different, but such differences are in all likelihood directly linked to the quality of the Freesurfer output given the data of a particular pulse-sequence. There is no reason to assume that results might be different with comparable accuracy of the surface reconstruction and subsequent cortical labeling.

For fitting and interpolating borders we used high-order polynomial models. Different fitting approaches might have been used, such as techniques based on Fourier transformation. The amount of mean fitting error was however small (0.43 mm) relative to the spatial resolution of most pulse sequences, or any other potential sources of positioning error e.g. those related to the intersubject variation in functional anatomy (>8 mm) (Mikl et al., 2008). Furthermore, the largest sources of error were those caused by situations where the assumption of a single y-coordinate for each x-coordinate was incorrect, and changing the fitting procedure cannot provide improvement. The chosen fitting procedure is thus in all likelihood close to optimal within the functional framework of the toolbox.

While the toolbox has a potential for broad application, there are a couple of inherent limitations to its usage.

1: A Cgrid requires the definition of 4 ROI-borders, which prevents the generation Cgrids of some specific ROIs or combination of ROIs as defined by one of the cortical labeling schemes. Note however that any portion of cortex can be covered by using alternative selection of ROI-borders.

2: The proposed approach is based on surface reconstruction of cortex, which basically limits its application to portions of cortex for which this can be generated accurately. Where the cortical sheath is damaged as a result of stroke or neurologic disease, the application will be more complicated.

3: Although for most Cgrid-schemes that we tested, the toolbox performed near perfectly, as estimated by visual inspection of results and the amount of error when fitting polynomials to the borders, for a number schemes there were occasional failures of the algorithm, inducing relatively large inaccuracies in the produced Cgrids. These included Cgrids covering the insula, medial orbitofrontal cortex, rostral anterior cingulate and the parahippocampus, as defined using the DK-atlas (Desikan et al., 2006). Several different causes were underlying these failures, including errors in the cortical annotation and the flattening, that resulted in islands and crossovers in the borders that defined the Cgrids. Other issues were more directly linked to the native functions within the toolbox, including errors in the estimation of start/end points of interpolated polynomials, and errors in the extrapolation of borders. This stresses the necessity for validating newly developed Cgrid schemes, using the optional outputs of the toolbox for visual inspection. Note also that most of the aforementioned issues can be avoided by alternative choices of borders that compose the Cgrid.

4: Cgrids can include extensive portions of cortex, but the toolbox so far does not support full brain coverage within a single Cgrid. While the registration framework proposed by Auzias et al. (Auzias et al., 2013) includes a whole brain rectangular representation, it does not offer the level of flexibility that is provided by the Cgrid toolbox. In addition, near full brain coverage can in principle be achieved by generating multiple Cgrids per subjects and combining results. Future developments might allow for more complete coverage of single Cgrids, achieved by using alternative custom build cortical labeling schemes or by adapting the algorithms so that Cgrids of different ROIs can be flexibly concatenated.

5: Multicomparison correction of statistical maps based on Cgrids can be complicated by non-uniform deformations across the Cgrids, which can in turn induce spatial variations in local smoothness of results. While classic Bonferroni correction for the number of Cgrid tiles will be a safe (but rather harsh) approach for avoiding false positives, such smoothness variations make cluster level inference e.g. through random field theory difficult to implement (Friston et al., 1996; Poline et al., 1997). Applying such methods on Cgrids will need to take variations in local smoothness into account, especially when the amount of spatial deformation induced in making the Cgrid is large.

In conclusion, the Cgrid approach allows to build rectangular representations of brain areas in order to facilitate spatial orientation and pattern recognition in addition to several other advantages. For most brain areas the algorithm works close to perfect, while for a small number of areas it encountered occasional problems. Future versions will aim at improved handling of situations that now result in errors and incorporating algorithms that allow concatenating different Cgrids, to cover larger portions of the brain.

Table 1: Parameters of the Cgrid schemes and results of the evaluation; 1: Name of the scheme;2: Names of the ROIs composing the subgrids;3: The FreeSurfer parcellation scheme used for the Cgrid, including DK (Desikan et al., 2006) and DKT (Desikan et al., 2006);4: x-resolution for each subgrid;5: y-resolution for each subgrid;6: The fitting approach used for estimating the x-coordinates, including 2 (2-polynomial approach) or 4 (4 polynomial approach);7: The kernel size used for smoothing the left and right border (mm FWHM);8: Mean±SD (across subjects) of the root mean square error of the border fits;9: Mean±SD (across subjects) of the maximum deviation between the border coordinates and the polynomial model;10: % of Cgrids affected by border extrapolation errors: ;11:% of Cgrids not generated because not all borders detected ;12:% of Cgrids affected by crossovers: ;13:% of Cgrids affected by islands;14:% of Cgrids affected by interpolated polynomials crossing the left or right border at multiple locations;15:% of all Cgrids affected by errors

