## Supplementary material for "Putting the brain in a box: a toolbox for creating Cartesian Geometric Representations with Isometric Dimensions (Cgrids)": Table 1

| **CGRID^1^** | **Subgrid^2^** | **Parcellation scheme^3^** | **x-dim^4^** | **y-dim^5^** | **Fitting approach^6^** | **Border Smoothing^7^** | **Mean Fit Error (mm)^8^** | **Worst Fit Error (mm)^9^** | **Error 1(%)^10^** | **Error 2(%)^11^** | **Error 3(%)^12^** | **Error 4(%)^13^** | **Error 5(%)^14^** | **Error All(%)^15^** |
| --- | --- | --- | --- | --- | --- | --- | --- | --- | --- | --- | --- | --- | --- | --- |
| Caudalmiddlefrontal | 1: Caudalmiddlefrontal | DK | 30 | 30 | 4 | NA | 0.22±0.06 | 1.27±0.62 | 0.00 | 0.00 | 0.00 | 0.00 | 0.00 | 0.00 |
| Cingulate | 1: Posteriorcingulate | DK | 42 | 21 | 2 | 4 | 0.33±0.14 | 1.49±0.87 | 0.00 | 0.00 | 0.00 | 0.00 | 0.00 | 0.00 |
|  | 2: Caudalanteriorcingulate |  | 42 | 21 | 2 | 4 | 0.20±0.10 | 0.89±0.68 | 0.00 | 0.00 | 0.00 | 0.00 | 0.00 | 0.00 |
|  | 3: Rostralanteriorcingulate |  | 42 | 21 | 2 | 4 | 0.61±0.30 | 3.05±1.35 | 0.00 | 0.00 | 1.67 | 0.00 | 20.00 | 21.67 |
| Insula | 1: Insula | DK | 90 | 30 | 4 | NA | 0.95±0.35 | 6.92±2.47 | 0.00 | 0.00 | 51.67 | 0.00 | 0.00 | 51.67 |
| Isthmuscingulate | 1: Isthmuscingulate | DK | 42 | 21 | 4 | NA | 0.32±0.09 | 2.54±1.08 | 0.00 | 0.00 | 0.00 | 0.00 | 0.00 | 0.00 |
| Language | 1: Parsopercularis | DKT | 38 | 18 | 2 | 2 | 0.17±0.03 | 0.58±0.16 | 0.00 | 0.00 | 0.00 | 0.00 | 0.00 | 0.00 |
|  | 2: Parstriangularis |  | 38 | 24 | 2 | 2 | 0.23±0.05 | 0.95±0.31 | 0.00 | 0.00 | 0.00 | 0.00 | 0.00 | 0.00 |
|  | 3: Parsorbitalis |  | 38 | 18 | 2 | 2 | 0.23±0.04 | 0.81±0.23 | 0.00 | 0.00 | 0.00 | 0.00 | 0.00 | 0.00 |
| Lateraloccipital left | 1: Lateraloccipital | DK | 60 | 30 | 4 | NA | 0.64±0.51 | 4.61±4.09 | 1.67 | 0.00 | 1.67 | 0.00 | 0.00 | 3.33 |
| Lateralorbitofrontal | 1: Lateralorbitofrontal | DK | 30 | 15 | 4 | NA | 0.54±0.21 | 4.36±2.15 | 8.33 | 0.00 | 0.00 | 0.00 | 0.00 | 8.33 |
| Medialorbitofrontal | 1: Medialorbitofrontal | DK | 60 | 30 | 2 | 4 | 1.01±0.51 | 5.32±2.92 | 16.67 | 3.33 | 0.00 | 0.00 | 1.67 | 21.67 |
| Paracentral | 1: Paracentral | DK | 36 | 36 | 4 | NA | 0.16±0.02 | 0.70±0.24 | 0.00 | 0.00 | 0.00 | 0.00 | 0.00 | 0.00 |
| Parahippicampal | 1: Parahippocampal+Entorhinal | DK | 45 | 15 | 2 | 3 | 0.86±0.62 | 4.12±2.68 | 8.33 | 1.67 | 1.67 | 3.33 | 16.67 | 31.67 |
| Parietal | 1: Supramarginal+Inferiorparietal | DK | 90 | 45 | 2 | 2 | 0.48±0.08 | 1.93±0.50 | 0.00 | 0.00 | 0.00 | 0.00 | 0.00 | 0.00 |
|  | 2: Superiorparietal |  | 90 | 45 | 2 | 2 | 0.51±0.09 | 2.16±0.69 | 0.00 | 0.00 | 0.00 | 0.00 | 0.00 | 0.00 |
| Precuneus | 1: Precuneus | DK | 60 | 30 | 4 | NA | 0.34±0.09 | 1.87±1.01 | 0.00 | 0.00 | 0.00 | 0.00 | 0.00 | 0.00 |
| Rostralmiddlefrontal | 1: Rostralmiddlefrontal | DK | 120 | 60 | 4 | NA | 0.46±0.14 | 2.73±1.30 | 1.67 | 0.00 | 0.00 | 0.00 | 0.00 | 1.67 |
| Sensorimotor | 1: Precentral | DK | 80 | 20 | 2 | 2 | 0.51±0.09 | 2.20±0.59 | 0.00 | 0.00 | 0.00 | 0.00 | 0.00 | 0.00 |
|  | 2: Postcentral |  | 80 | 20 | 2 | 2 | 0.41±0.07 | 1.59±0.49 | 0.00 | 0.00 | 0.00 | 0.00 | 0.00 | 0.00 |
| Superiorfrontal | 1: Superiorfrontal | DK | 80 | 40 | 2 | 2 | 0.76±0.21 | 4.49±1.74 | 1.67 | 0.00 | 0.00 | 0.00 | 0.00 | 1.67 |
| Temporal | 1: Fusiform | DK | 96 | 24 | 2 | 2 | 0.36±0.10 | 1.47±0.67 | 0.00 | 0.00 | 0.00 | 0.00 | 0.00 | 0.00 |
|  | 2: Inferiortemporal |  | 96 | 24 | 2 | 2 | 0.35±0.06 | 1.31±0.35 | 0.00 | 0.00 | 0.00 | 0.00 | 0.00 | 0.00 |
|  | 3: Middle/Superior/Transverse-temporal+Bankssts |  | 96 | 72 | 2 | 2 | 0.38±0.07 | 1.61±0.46 | 0.00 | 0.00 | 0.00 | 0.00 | 1.67 | 1.67 |
| Visual | 1: Lingual | DK | 60 | 30 | 2 | 2 | 0.25±0.08 | 1.07±0.43 | 0.00 | 0.00 | 0.00 | 0.00 | 0.00 | 0.00 |
|  | 2: Pericalcerine |  | 60 | 15 | 2 | 2 | 0.30±0.11 | 1.14±0.44 | 0.00 | 0.00 | 0.00 | 0.00 | 1.67 | 1.67 |
|  | 3: Cuneus |  | 60 | 15 | 2 | 2 | 0.21±0.06 | 0.80±0.26 | 0.00 | 0.00 | 0.00 | 0.00 | 0.00 | 0.00 |
|  |  |  |  |  |  | **Mean** | 0.44 | 2.30 | 1.42 | 0.19 | 2.10 | 0.12 | 1.54 | 5.37 |
